# Multi-modal Spatial-modality Attentive Fusion for Studying Neuropsychiatric Disorders

**DOI:** 10.1101/2024.06.09.598091

**Authors:** Md Abdur Rahaman, Yash Garg, Armin Iraji, Zening Fu, Peter Kochunov, L. Elliot Hong, Theo G. M. Van Erp, Adrian Preda, Jiayu Chen, Vince Calhoun

## Abstract

Multi-modal learning has emerged as a powerful technique that leverages diverse data sources to enhance learning and decision-making processes. Adapting this approach to analyzing data collected from different biological domains is intuitive, especially for studying neuropsychiatric disorders. A complex neuropsychiatric disorder like schizophrenia (SZ) can affect multiple aspects of the brain and biologies. These biological sources each present distinct yet correlated expressions of subjects’ underlying physiological processes. Joint learning from these data sources can improve our understanding of the disorder. However, combining these biological sources is challenging for several reasons: (i) observations are domains-specific, leading to data being represented in dissimilar subspaces, and (ii) fused data is often noisy and high-dimensional, making it challenging to identify relevant information. To address these challenges, we propose a multi-modal artificial intelligence (AI) model with a novel fusion module inspired by a bottleneck attention module (BAM). We use deep neural networks (DNN) to learn latent space representations of the input streams. Next, we introduce a two-dimensional (spatio-modality) attention module to regulate the intermediate fusion for SZ classification. We implement spatial attention via a dilated convolutional neural network that creates large receptive fields for extracting significant contextual patterns. The resulting joint learning framework maximizes complementarity allowing us to explore the correspondence among the modalities. We test our model on a multi-modal imaging-genetic dataset and achieve an SZ prediction accuracy of 94.10% (P < 0.0001), outperforming state-of-the-art unimodal and multi-modal models for the task. Moreover, the model provides inherent interpretability that helps identify concepts significant for the neural network’s decision and explains the underlying physiopathology of the disorder. Results also show that functional connectivity among subcortical, sensorimotor, and cognitive control domains plays an important role in characterizing SZ. Analysis of the spatio-modality attention scores suggests that structural components like the supplementary motor area, caudate, and insula play a significant role in SZ. Biclustering the attention scores discover a multi-modal cluster that includes genes CSMD1, ATK3, MOB4, and HSPE1, all of which have been identified as relevant to schizophrenia. In summary, feature attribution appears to be especially useful for probing the transient and confined but decisive patterns of complex disorders, and it shows promise for extensive applicability in future studies.

## I. Introduction

**O**ur brain processes multi-modal signals from the outer world to understand real events and respond accordingly [1-5]. Leveraging information from diverse sources allows for a better understanding of a given phenomenon. Likewise, multi-modal learning enables researchers and practitioners to benefit from the unique strengths of each modality, leading to improved performance, enhanced accuracy, and richer insights [6, 7]. Applications of multi-modal learning models are wide-ranging and include fields such as computer vision [8-10], healthcare [11-13], and speech recognition [14, 15], among others [16]. Moreover, the potential applications of multi-modal learning in studying mental disorders are vast and continue to expand [17-22]. The rationale lies in multiple biological domains that are affected by the underlying physical conditions. Therefore, maximizing the complementarity among these sources can potentially lead to a more comprehensive understanding of the data.

Research indicates that useful biological information is encoded across different sources. Neuroimaging provides structural and functional information about the brain through various imaging modalities [17, 22]. Genetic data provide another source of information regarding disease-related aberrations [23, 24]. The integration of data from different modalities and sources can offer a richer and more nuanced understanding of complex mental disorders. Data from multiple modalities are not mutually exclusive but complement each other in describing brain processes [22]. Particularly, neuroimaging modalities, when combined can achieve enhanced temporal and spatial resolution and bridge the gap between physiological and cognitive representations [25]. Hence, multi-modal learning frameworks have emerged as effective tools for analyzing data from multiple sources, including neuroimaging [17, 20, 26, 27] and genetic data [18, 28]. Past research has demonstrated significant correlations between structural and functional changes in the brain and mental disorders [29, 30]. Moreover, existing scientific literature points to a promising area of exploration: the correlation between genetic variants and neural activity concerning neuropsychiatric disease-related degeneration [23, 24, 31]. Such a disorder Schizophrenia (SZ) is genetically complex and affects the brain’s structure and function [18, 32, 33]. Nevertheless, significant challenges arise in the joint analysis of genetic, structural, and functional data, as they often carry information at different scales and formats. A perceptive fusion module is necessary for enhancing the model’s performance as it ensures the judicious use of the most informative sources in subsequent tasks. A bottleneck strategy might facilitate the integration of these diverse knowledge domains, overcoming the obstacles posed by differing data scales and formats. The bottleneck in a neural network is a layer with fewer neurons than the layer below or above it [34, 35]. It helps to learn representations better and emphasize salient features for the target variable. Thus, our intuition is to operate fusion in this layer with bottleneck attention.

Research has provided a thorough classification of fusion methodologies [36, 37]. These fusion strategies are broadly divided into three subcategories: early fusion, mid/intermediate fusion, and late fusion, depending on the phase of the model where the integration takes place. The mid-fusion approach is particularly notable due to its wide range of applications. In this scheme, fusion occurs after feature extraction from different input modes [36]. However, integrating multiple sources in the intermediate phase presents a challenge, as differences in data types, collection methods, scales, and preprocessing can lead to inconsistencies in data representations. To address this issue, modality-specific neural networks are used to first learn the hidden space representations of the inputs. These networks translate the multi-modal inputs into a uniform embedding space, after which the mid-fusion module is used for the integration [37, 38]. It is important to note that even when employing latent space fusion, the resulting fused feature map can be high-dimensional, with each unit’s contribution to the subsequent task potentially varying in importance. Consequently, simply connecting the fused tensor to the predictor may limit the model’s performance. This problem is especially pronounced when data modalities are unevenly informative about the downstream task, a common occurrence in medical data collection. Neuroimaging and genomics datasets often contain weak descriptors of the underlying biology for a few samples. Therefore, the fusion module needs to enhance the synergies between these data sources. Previous research has proposed various embedding fusion techniques, including attention-based [39, 40], multiplicative combination layer [41], and transformer [42]. An early adaptation of the bottleneck attention module (BAM) [35] known as the mBAM [43] examined both spatial and modality dimensions. It employed a fully connected (FC) layer with compression to learn spatial attention from the fused two-dimensional tensor.

In this study, we present a fusion module called spatio-modality fusion using bottleneck attention which carefully examines the amalgamation. We use dilated convolutional [44] methods to learn the contextual pattern. The module explores spatial dimensions using a convolutional neural network (CNN) [45] augmented by a larger receptive field (RF), a feature known as dilated convolution. Using dilated convolutions significantly expands the receptive field, facilitating the collection of contextual patterns from the combined data [35]. These contextual patterns play a crucial role in the downstream tasks. Our study shows that the dilated convolutional layer performs better than the FC layers (Fig. 3). In earlier studies, the FC layers lost considerable spatial information while downsampling from the combined tensor to a one-dimensional vector [43]. Furthermore, the subsequent attention operation performed on the compressed version produced a low-dimensional (vector) mask. Our method, on the other hand, implements spatial attention on the original version of the fused tensor, generating a two-dimensional attention mask. Simultaneously, the module learns to apply attention across the modalities to select the best source, and in spatial dimensions to mask the compressed feature vectors, fostering a richer knowledge extraction. The attention mask delineates the significance of each feature on classification, in our case identifying schizophrenia. Therefore, the mask can also be utilized to generate other analytics of the test samples e.g., subgrouping based on disease relevance.

The modalities we use for the classification are structural and functional neuroimaging data and genome-wide polymorphism data. We test the model on a dataset comprising 437 subjects, including individuals with schizophrenia (162) and controls (275), intending to classify SZ. Our proposed method produces a multi-dimensional attention mask to elucidate the model’s decisions and underlying neurobiological basis. This attention mask encodes the relative importance of each modality and spatial significance. The spatio-modality attention identifies sMRI components – such as the supplementary motor area (SMA), left insula, caudate, and temporal pole – of high importance for detecting schizophrenia (Figure 4). The attention scores on static functional connectivity suggest that several connections among the sensorimotor, subcortical, and cognitive control are particularly salient in schizophrenia. Biclustering the attention scores from three modalities discovers a multi-modal cluster that includes a subset of relevant structural components, functional connections, and genes. The implicated genes, CSMD1, ATK3, MOB4, and HSPE1, have been previously recognized as relevant to schizophrenia. The primary features of our method are:

1. A fusion module capable of encoding both modality and spatial attention.
2. A dilated convolution to extract spatial patterns with a large receptive field.
3. A two-dimensional spatio-modality attention mask, enabling further data analytics such as subgroup identification.
4. A self-explainable model via the attention scores for modality and contextual dimension.

## II. Data Preprocessing

### A. Structural MRI

We preprocessed structural magnetic resonance imaging (sMRI) scans using statistical parametric mapping (SPM12, http://www.fil.ion.ucl.ac.uk/spm/). The preprocessing steps include unified segmentation and normalization of sMRI scans into gray matter, white matter, and CSF. During the segmentation step, we use a modulated normalization algorithm to generate gray matter volume (GMV). Following this, we use a Gaussian kernel with a full width at half maximum (FWHM) of 6 mm to smooth the gray matter densities.

### B. Functional MRI

We use the SPM12 toolbox for preprocessing functional MRI data. To ensure steady-state magnetization, we remove the first five time points of the fMRI. Next, we carry out rigid body motion correction using the INRI-Align robust M-estimation approach and apply the slice-timing correction. We then spatially normalized fMRI images into the Montreal Neurological Institute (MNI) standard space, using an echo-planar imaging (EPI) template, and resampled to 3 × 3 × 3 mm^3^ isotropic voxels. The images are then smoothed with a Gaussian kernel with FWHM = 6 mm, similar to the sMRI data.

### C. Genomics

The preprocessing steps for the genetic data are described in our prior work [46, 47]. There are several standard preprocessing tools for genomics data, and we use Plink [48] for both preprocessing and imputation. Linkage disequilibrium (LD) pruning was administered at r^2^ < 0.9. We use the Psychiatric Genomics Consortium (PGC) for SZ suggested GWAS [48] score to select the features. The analysis selects 1280 SNPs distributed across 108 risk loci. The PGC study [49] reveals these SNPs express statistically significant associations with schizophrenia at p < 1×10^−4^. The datasets verify the test-retest experiment for MRI acquisitions by recording multiple repetitive scans for each subject and also account for the stable MRI signal.

More details about preprocessing and quality control are described in these studies [50, 51].

## III. Our proposed Multi-modal architecture

Figure 1 demonstrates our proposed architecture for multi-modal learning. The model incorporates three modality-specific neural networks (NN), referred to as subnetworks, which are used for learning the latent space embedding from each input source. These neural networks are selected based on their effectiveness given the type of input data, and we empirically validate their efficacy. The subnetworks conduct dimensionality reduction and account for missing entries and other discrepancies to effectively learn the representations from each incoming modality. The latent embeddings from all modalities are concatenated into a multi-dimensional tensor. This tensor goes through a novel spatio-modality attention-based fusion module, which is described in Figure 2. Once this is completed, the two-dimensionally attended fused embedding is used as input to a multi-layer perceptron (MLP). This is followed by a SoftMax layer for classification. This approach allows us to extract rich features from multiple modalities and harness the power of neural networks to classify complex data effectively.

**Fig. 1.**
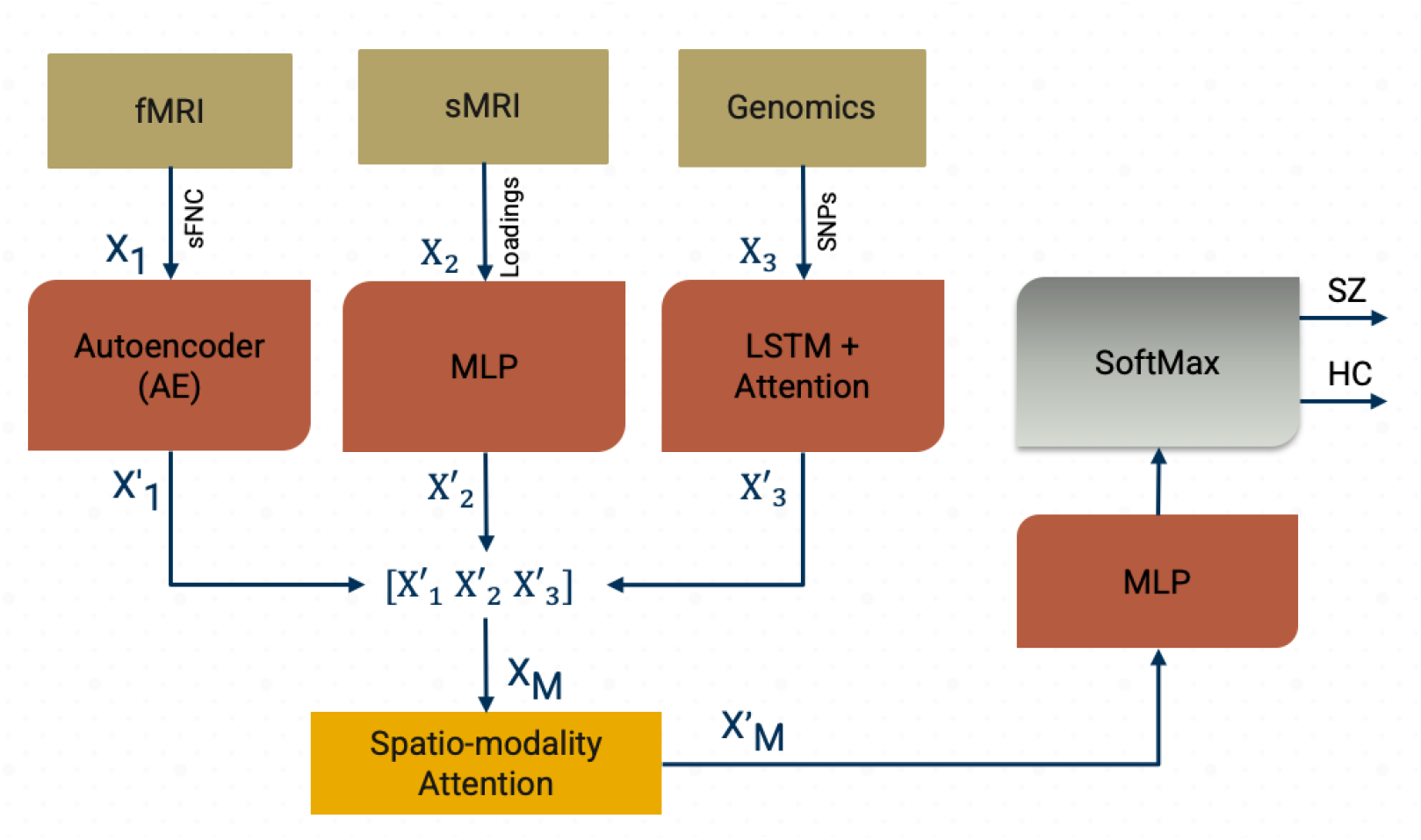
Our proposed multi-modal framework with spatio-modality temporal attention. These models incorporate three major processing steps: imaging-genomics preprocessing and dimensionality reduction, neural subnetworks for learning latent space embedding, and the predictor. We run group ICA on sMRI and fMRI data. We generate static functional network connectivity (sFNC) among the independent component networks (ICNs) extracted from fMRI decomposition. We select the ICA loading parameter as the input feature for sMRI modality. The GWAS-based genetic variable selection is carried out for genetic modality. The subnetworks are deep neural networks for learning the modality-wise representation. Subnetwork 1 is an autoencoder (AE) for learning sFNC, subnetwork 2 is a multi-layered perceptron (MLP) for sMRI loadings, and subnetwork 3 is a bi-directional long short-term memory (LSTM) unit with attention for learning SNPs. The combined embedding is attended in spatial and modality direction and sent through an MLP followed by a SoftMax layer for the classification. The architecture is jointly trained using an Adam optimizer.

**Fig. 2.**
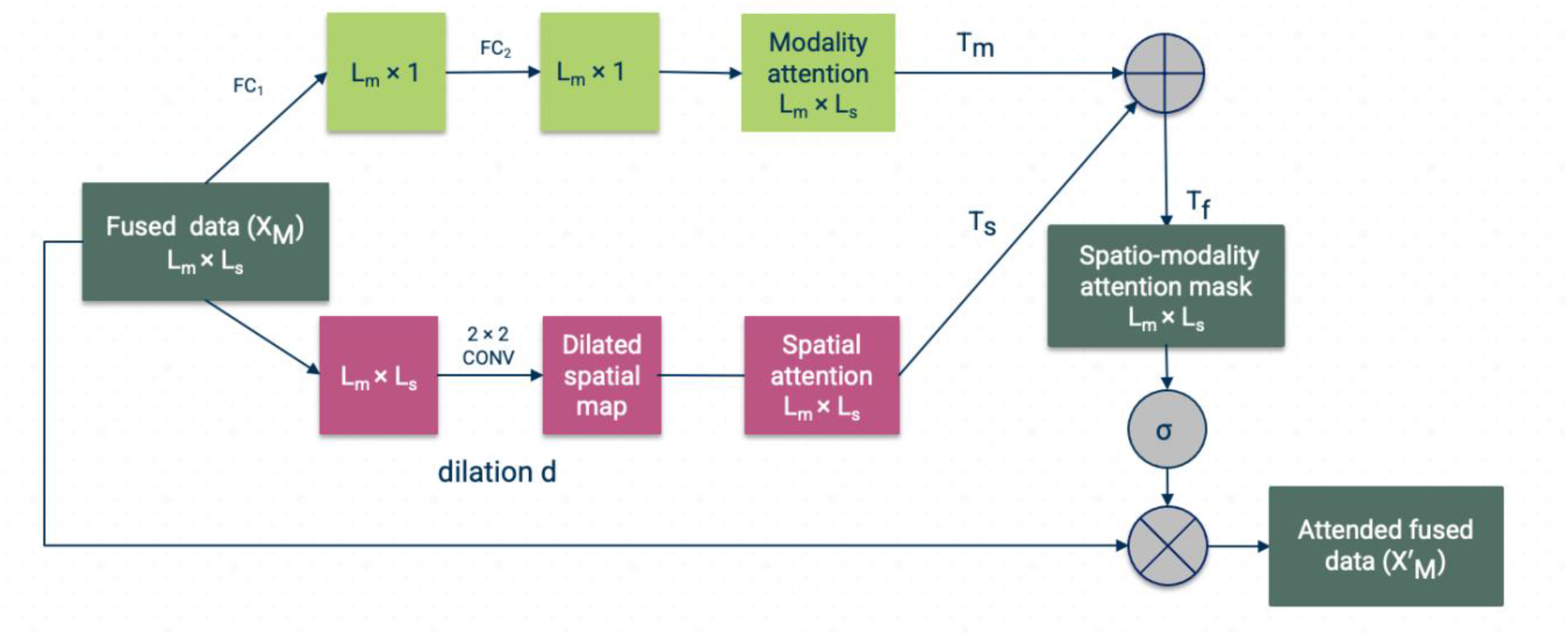
Our proposed spatio-modality attention module for multi-modal fusion. The concatenated embeddings are sent through two different branches. i) The modality branch that learns the cross-modality interactions and mounts it into T_m_ attention mask, ii) the spatial branch (T_s_) captures the relevant context from each biological site. These two masks are merged into a final attention mask T_f_. The spatial branch uses dilated convolution for learning the contextual understanding of the multi-modal tensor and fully connected layer for modality attention.

### A. Input features quantification

This phase of the framework includes a few processing units for refining and performing the initial decomposition of the data collected from multiple biological domains. We execute a fully automated spatially constrained group ICA (GICA) using the Group ICA of fMRI Toolbox (GIFT) available at this link: (http://trendscenter.org/software/gift) [52] and the NeuroMark [46] template on the combined subjects from three datasets, which include FBIRN [53], COBRE [26], and MPRC [46]. The selection of independent component networks (ICNs) in this study is based on the NeuroMark template. We selected 53 pairs of ICNs and arranged them into seven functional networks based on prior anatomical and functional knowledge fields [50, 51]. The total number of connectivity networks extracted was 53, covering the whole brain. These intrinsic connectivity networks (ICNs) were distributed into seven functional domains [50, 54]: sub-cortical (SC), auditory (AD), sensorimotor (SM), visual (VS), cognitive-control (CC), default-mode (DM), and cerebellar domain (CB). After estimating subject-specific networks, we compute the subject-wise static functional network connectivity (sFNC). The square matrix (53×53) represents the Pearson correlation between the time course (TCs) of ICNs. We vectorize the sFNC matrix using the upper diagonal entries to ease the encoder’s training process. A similar approach was used for structural MRI source-based morphometry (ICA on gray matter maps) [55, 56], resulting in 30 structural components along with their loading values. The provided ICA priors are estimated from the 6500 subjects used in the referred study [57]. The ICA on sMRI yields individual-level structural networks and corresponding loadings. For the sMRI modality, we used the loadings of the structural networks as the structural features. The third modality includes the set of SNPs selected from genomic data based on GWAS-significant schizophrenia risk SNPs identified by the large PGC study.

### B. Deep neural networks (DNNs)

We employ distinct DNNs for extracting modality-wise features. The design choices are explained in the following subsections, and their efficacy is tested on the dataset. We use an autoencoder (AE) [58] to learn the representation of sFNC. It follows an encoder and decoder architecture, with each submodule consisting of five linear layers. The encoder compresses the input and generates a compressed embedding. The decoder reconstructs the input from the encoded features map. The loss function computes the reconstruction loss as the mean squared error (MSE). We use the sFNC matrix as an input from the fMRI modality and employ an AE with rectified linear unit (ReLU) activation for learning the representations. AE is effective for learning semantic meanings and compressed abstraction of input data with successful applications in diverse fields of study [59, 60]. We use Xavier initialization [61] from Pytorch to set the initial values of the layers in the network. We applied a dropout strategy with a probability p of 0.2 to minimize overfitting. Overfitting refers to a situation where a model performs well on the training data but fails to generalize to unseen data. Dropout is a regularization technique that helps to mitigate this issue by randomly ignoring a subset of features during training, reducing the complexity of the model and promoting generalization. For the sMRI subnetwork, we use the loading parameters from group ICA. The feature vector has a length of 30, where each value represents the expression level of a subject on a group structural independent component (GsIC). In general, loadings are just betas/coefficients of the variance mixture linear equations [62]. The loading value corresponds to how much variance a subject contributes to a given group component. Since the loadings are the vector of discrete values; for simplicity, we deployed a multi-layer perceptron (MLP) to efficiently learn the sMRI features. An MLP is a type of artificial neural network that consists of multiple layers of nodes in a directed graph, with each layer fully connected to the next one. It can model complex, non-linear relationships between input and output data, making it a suitable choice for feature extraction in this context. Each layer of the MLP takes the output of the previous layer (or the input data for the first layer), applies a set of weights (learned during training), and then applies an activation function, producing the output for that layer. By adjusting the weights during training, the MLP can learn to extract salient features from the input data that are informative for the task at hand, which in this case, is the classification of schizophrenia. The subnetwork is fully connected since the data dimension (30) is lower than the other modalities. In the final layer of our network, we dilated the embeddings to 100 to ensure consistency with the size of latent features derived from different modalities. We implemented a bi-directional long short-term memory (bi-LSTM) network with an attention mechanism [63] to extract features from SNPs. The use of a bi-LSTM was informed by its anticipated ability to capture contextual information from a sequence [64, 65]. We assume that neighboring SNPs may show linkage disequilibrium with one other, potentially forming a neighborhood substructure. The bi-LSTM is designed to capture the localized semantics of genomic data, which might help differentiate between cases of the disorder and controls. In addition, the attention scores signify contributing neighbors to the context, further enhancing the deep neural network’s discriminative power.

### C. Spatio-modality Attention

We propose a fusion module that takes a multi-modal joint embedding and probes the concatenated tensor in both modality and spatial dimensions. The architecture is demonstrated in Figure 2. The architecture is inspired by the BAM [35] which has been implemented for learning channels and spatial attention in classification and prediction tasks [66-68]. We adapt the module for fusing different modalities. The module is integrated with a mid/intermediate fusion [69] in a multi-modal learning framework. In mid-fusion, the modality-specific DNNs generate compressed representations of input streams (bottleneck layers) from each source. Our intuition is to run two-dimensional attention on these bottleneck layers. Our module uses two simultaneous attention branches that forage significant bits from the fused tensor. These branches can be treated as two distinct scoring functions across modality and spatial dimensions. We use fully connected (FC) linear layers in modality dimensions to learn the attention weights per modality. Fully Connected layer (FC), is a type of layer used in neural networks where all neurons in the previous layer are connected to the neurons in the next layer. The FC operation is defined in the following equation (2).

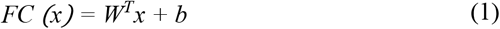

where *W ∈* ℝ^*N×1*^, *b ∈* ℝ^*N×1*^ is the weight matrix and bias respectively and *N* is the input dimension. *x* is the input to the FC operation, in the context of equation (1), and *b* is the bias vector in the fully connected layer.

For another branch, the module employs dilated convolution layers for extracting the relevant spatial patterns. The dilated convolutional filters arbitrate a large receptive field for collecting contextual information. In our study, we are using only three modalities, so we skip the reduction. However, the reduction ratio in modality direction could potentially help in combining a large number of modalities in further studies.

#### 1) Modality attention scoring (T_m_)

The modality attention branch assigns scores to the modality dimension to signify the most informative and discriminative source for the downstream task. The modality attentions mask (*T*_*m*_) gathers importance scores that characterize the contribution of input modality to the classification of SZ. Figure 2 illustrates the attention-weighting architecture. An FC linear layer (*FC*_*1*_) is applied to the input tensor, *X*_*M*_ (L_m_ × L_s_) to reduce the dimension to the number of modalities (L_m_). Here, L_s_ is defined as the latent embedding size consistent across the modalities. The compressed data is then passed through another FC layer (*FC*_*2*_) to compute the attention weights for each modality. We also use batch normalization (*BN*) [70] to adjust the scale with the spatial branch. Moreover, *BN* improves the speed, performance, and stability of the neural networks. Then, the tensor is expanded to the shape of the input tensor L_m_ × L_s_. We can formulate the operation as follows,

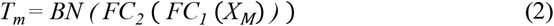

Here, *Tm* refers to the transformed tensor after the application of batch normalization and two fully connected layers.

#### 2) Spatial attention branch (Ts)

We opt for capturing the context from the aggregated multi-modal embedding tensor, which extracts contextual information in the form of spatial patterns. For spatial attention, the submodule uses a convolutional neural network. We use dilated convolution [44] to create a large receptive field that assists in apprehending the context. The dilated convolution inflates the kernel by inserting holes between the kernel elements [44]. An additional parameter, dilation ratio (*d*), indicates the extent to which the kernel is widened. There are usually *d*-1 spaces inserted between kernel entries. We use an empirically validated convolutional filter of dimension 2 × 2 with a dilation ratio (*d*) = 2 on the combined tensor. Figure 2 shows the dilated convolution for spatial significance. The branch weighs the fused data for identifying salient loci relevant to downstream tasks. The spatial attention mask is also expanded to a shape of L_m_ × L_s_. Equation 3 is the operation for spatial attention and illustrates the overall processing in the spatial branch.

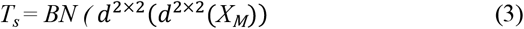

Here, *T*_*s*_ represents the spatial attention mask. The operation helps to create a large receptive field that captures the context or spatial pattern in the input data. We merge the spatial attention mask, *T*_*s*_ and the modality attention mask, *T*_*m*_ to generate the final mask, *T*_*f*_ (see equations (4)).

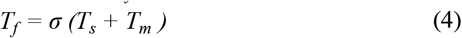

The operation in equation (4) is an element-wise summation between *T*_*s*_ and *T*_*m*_ to compute the final fused attention mask, *T*_*f*_. We apply a sigmoid function (*σ*) to bind the values of *T*_*f*_ between 0 to 1. The sigmoid function is commonly used in neural networks to introduce non-linearity and to map any input to a value between 0 and 1, making it especially suitable for models where we have to predict probabilities.

Next, we use element-wise multiplication operation (⊗) to fuse the attention mask *T*_*f*_ and the input tensor *X* (equation (5)).

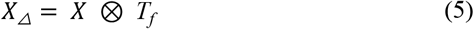

This operation highlights the important features in the input tensor according to the attention mask. The result is a new tensor matrix, which emphasizes the regions of the input that are most informative for the task at hand. We use *X*_*△*_ as the input to the MLP followed by a SoftMax layer to classify the samples (Figure 1). Together, these operations help to guide the model to focus on the most informative features across different modalities and spatial locations when performing the downstream task, such as classification or regression.

### D. Multi-modal joint training

We split the dataset into two primary categories: 80% for training and 20% for evaluation. The evaluation data is then further divided equally into testing and validation sets. We train the joint model for 450 training epochs and validate its performance on the validation set. For the training phase, we use the *Adam* optimizer from Pytorch with a learning rate (LR) of 10^−4^ and a batch size of 32. The loss terms include mean square error (MSE) for the autoencoder subnetwork reconstruction loss and binary cross-entropy for the model’s classification loss. We implement early stopping to balance the training and validation loss, which eventually regularizes the model. Our training scheme optimizes for accuracy and saves the best-performing model. For the validation phase, we used k-fold cross validation [71] technique for k = 10. The cross-validation method randomly divides the datasets into 10 equal partitions and uses 9 of them to train the model and the remaining one for testing the performance. The technique interchanges the training and testing set and repeats the process ten times. In the testing phase, we utilize the best-performing model saved from the validation phase and apply it to the testing samples. Our performance metrics, which include accuracy, precision, recall, and F1 score, are computed in this phase. We report the final performance on the held-out test dataset. To empirically validate the results and other architectural specifications, we run the model 50 times and report average results across these repetitions with corresponding standard deviation. We also introduced multiple random initializations [72] of the neural network to ensure the agnosticism of the model to different initial conditions. The resulting metrics in Table 1 are summarized across distinct intializations. The joint training of all three modalities allows the sources to interact and helps modality-specific subnetworks to optimally converge by leveraging learning from the other subnetworks. Moreover, we implement a multi-modal regularization technique for the completeness of the experiments. This technique aims to eliminate bias by maximizing the functional entropies [73]. We designed our implementation based on the existing regularizer and utilities.

**Table 1.** Performance comparison of our proposed model with several baselines (unimodal, bi-modal, and tri-modal) and the state-of-the-art models for a similar task. The performance metrics are presented as (mean ± standard deviation) across 50 repetitive runs.

| Models | Modalities | Data | Accuracy | Precision | Recall | F1 score |
| --- | --- | --- | --- | --- | --- | --- |
| Autoencoder | Unimodal | fMRI | $0.811 \pm 0.15$ | $0.587 \pm 0.11$ | $0.451 \pm 0.05$ | $0.510 \pm 0.11$ |
| Multi-layer perceptron | | sMRI | $0.782 \pm 0.13$ | $0.439 \pm 0.12$ | $0.477 \pm 0.04$ | $0.443 \pm 0.09$ |
| bi-LSTM with attention | | SNPs | $0.673 \pm 0.15$ | $0.338 \pm 0.13$ | $0.498 \pm 0.04$ | $0.403 \pm 0.08$ |
| Mid fusion | Bi-modal | sMRI + fMRI | $0.835 \pm 0.14$ | $0.577 \pm 0.09$ | $0.478 \pm 0.03$ | $0.523 \pm 0.01$ |
| Mid fusion | | SNPs + sMRI | $0.784 \pm 0.18$ | $0.397 \pm 0.08$ | $0.455 \pm 0.03$ | $0.424 \pm 0.15$ |
| Mid fusion | | SNPs + fMRI | $0.813 \pm 0.14$ | $0.501 \pm 0.05$ | $0.443 \pm 0.03$ | $0.470 \pm 0.01$ |
| Early fusion | Multi-modal | sMRI + fMRI + SNPs | $0.701 \pm 0.11$ | $0.575 \pm 0.12$ | $0.460 \pm 0.03$ | $0.511 \pm 0.02$ |
| Late fusion | | sMRI + fMRI + SNPs | $0.781 \pm 0.08$ | $0.615 \pm 0.12$ | $0.400 \pm 0.04$ | $0.485 \pm 0.03$ |
| Mid-fusion [18] | | sMRI + fMRI + SNPs | $0.876 \pm 0.06$ | $0.640 \pm 0.07$ | $0.501 \pm 0.04$ | $0.562 \pm 0.04$ |
| Mid-fusion with attention [19] | | sMRI + fMRI + SNPs | $0.921 \pm 0.02$ | $0.798 \pm 0.07$ | $0.522 \pm 0.04$ | $0.631 \pm 0.05$ |
| Mid-fusion with mBAM [43] | | sMRI + fMRI + SNPs | $0.932 \pm 0.04$ | $0.903 \pm 0.06$ | $0.561 \pm 0.05$ | $0.692 \pm 0.04$ |
| Spatio-modality mid fusion | | sMRI + fMRI + SNPs | $0.941 \pm 0.05$ | $0.829 \pm 0.06$ | $0.601 \pm 0.06$ | $0.697 \pm 0.05$ |

## IV. Experimental Results & Discussion

We analyzed three datasets, COBRE [74], fBIRN [53], and MPRC [46]. The merged dataset consists of 437 subjects, with 275 healthy controls (HC) and 162 subjects with schizophrenia (SZ). The performance of our proposed model is benchmarked against several other models, including unimodal and multi-modal baselines using imaging-genetics datasets. The performance comparison is detailed in Table 1. In early fusion, input data from all modalities are merged at the first step and the resultant vector is then processed through a neural network. For late fusion, we use different networks for distinct modalities. Each network classifies a sample based on the data it receives and a max voting scheme [75] is used to determine the final label for each instance. Our proposed spatio-modality attentive fusion model outperforms the comparing methods (Table 1), achieving an accuracy of 94.1% in differentiating schizophrenia. Other performance metrics are either superior or comparable to the benchmark models. Since our dataset is slightly imbalanced, we used the harmonic mean of precision and recall given by the F1 score [76] for the classification performance evaluation. Our proposed model achieves an F1 of 0.697 with a reasonable balance between precision and recalls expected in a biological population. Substantial performance improvements are achieved when the bottleneck attentive module (BAM) is used for fusion (as shown in the last two rows of Table 1), highlighting the utility of BAM in multi-modal fusion. The performance of the model using only genetic data is consistently low. This suggests that genomic data is solely an insufficient descriptor of phenomena i.e., discriminating schizophrenia. However, the noteworthy observation is the ability of the genomic data to complement other modalities in multi-modal settings, especially when spatio-modality fusion is employed. This underscores the capacity of the fusion module to leverage the contributions from various modalities with greater precision. Our model can suppress the features from less informative sources while prioritizing those from more informative ones, a desirable characteristic to effectively learn the multi-modal representation of input data.

### A. Reproducibility and reliability of the results

Reproducibility is a fundamental aspect that speaks to the reliability and validity of the computational findings [77, 78]. The crucial medical research domain such as neuroimaging signifies the utility of replication most. It allows for the independent verification of results across analytical frameworks, enhancing the adaptability of vital neurobiological outcomes. In this study, we evaluate the reproducibility of the model’s performance on the neuroimaging population by employing cross-validation strategies to assess the generalizability and stability of the results across different subsets of the data. We combine three datasets in this study COBRE, fBIRN, and MPRC, and preproces them under the same imaging protocols. Rationally, we create three parts of the combined dataset and we check the reproducibility of each subpopulation separately. For recording performance on each dataset, we train the model on two other datasets. For instance, COBRE performance is computed using the COBRE sample as held-out test data, and the model is trained on only samples from fBIRN and MPRC. Additionally, these experiments aid in eradicating the acquisition bias and explore the model’s efficacy on a smaller population. Furthermore, we generate two random splits of the data for the completeness of the experiments. It utilizes randomization training/validation and testing cohort selection. In the first random split (RDS 1), we select 40% of held-out data for testing and the remaining for training and validation. In the second one (RDS 2), we randomly select another 30 % for the held-out and the remaining for training and validation. Table 2 shows the model performance on these diverse randomizations of the dataset to provide a comprehensive understanding of the efficiency of such a framework in disease prediction. We observe the performances are reasonably reproducible from different settings of input data. Three parts of the combined dataset yield comparable results with the final performance of the proposed framework. Two random splits also exhibit proportionate metrics validating the reliability of the model in the downstream task which is classification in our case. However, with more data for training (RSD 2 in Table 2), the model achieves slightly better performance – standard in data-driven approaches. We also validate the extracted salient attributes of the data contributing to the schizophrenia characterization across these splits. The spatio-modality attention-based feature analysis in the following subsection is carried out on the summarized set of features stable through independent segments of the dataset To evaluate the impact of dilation on contextual learning, we conducted experiments using distinct dilatation d values. These also contribute to the verification of the model’s robustness on specific choice configurations. Figure 3 presents the results for three performance metrics: accuracy, precision, and F1-score. The bar graph showing the evaluation metrics also includes the confidence interval. It indicates a lower and upper bound of the data that describes a range or a corridor in which 95% of the predictions would fall given the actual true value. We employed dilated convolution to extract contextual information. The dilation rate d = 1 represents the standard convolution, while d greater than 1 denotes the dilated version. The model demonstrates superior performance with a dilation value of 2 and experiences a performance drop for higher values. These findings suggest that dilation helps in extracting contextual information through a larger receptive field and is beneficial for accurate prediction. Nevertheless, because our multi-modal fused tensor has limited dimensions, larger dilation might skip important transient patterns in the data by adding more holes in the receptive field. Due to the limited dimensions of the concatenated tensor, we are unable to explore the sensitivity beyond a dilation ratio of 3.

**Table 2.** Reproducible performance of the proposed method for diverse subsets of the data. The first three rows represent the performance on the independent dataset and the bottom two rows stand for random splits.

| Dataset/Split | Training & validation | Held-out test data | Accuracy | Precision | Recall | F1-Score |
| --- | --- | --- | --- | --- | --- | --- |
| COBRE | fBIRN + MPRC | COBRE | $0.920 \pm 0.08$ | $0.802 \pm 0.07$ | $0.575 \pm 0.05$ | $0.68 \pm 0.06$ |
| fBIRN | COBRE + MPRC | fBIRN | $0.931 \pm 0.09$ | $0.801 \pm 0.06$ | $0.586 \pm 0.05$ | $0.67 \pm 0.05$ |
| MPRC | COBRE + fBIRN | MPRC | $0.901 \pm 0.12$ | $0.787 \pm 0.09$ | $0.589 \pm 0.07$ | $0.674 \pm 0.07$ |
| RDS 1 | 60% of total data | 40% of total data | $0.932 \pm 0.08$ | $0.801 \pm 0.01$ | $0.608 \pm 0.05$ | $0.691 \pm 0.03$ |
| RDS 2 | 70% of total data | 30% of total data | $0.937 \pm 0.07$ | $0.854 \pm 0.03$ | $0.583 \pm 0.04$ | $0.693 \pm 0.03$ |

**Fig. 3.**
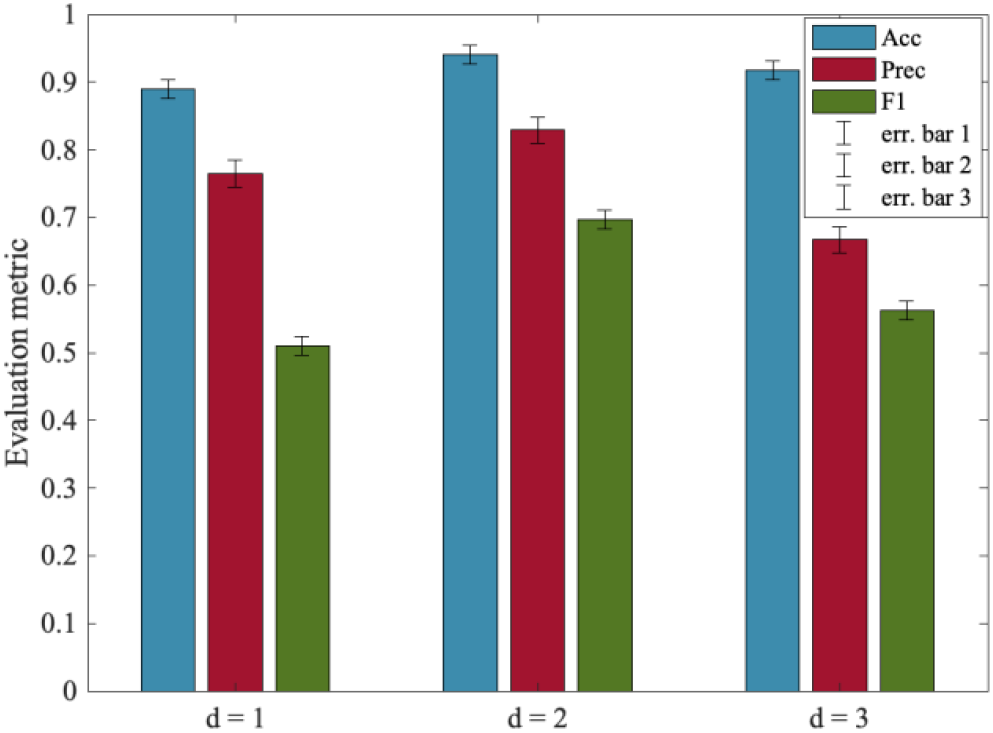
Model’s performance for various dilation rates (d). The error bars stand for the confidence interval for the metrics. The dilation rate indicates the expansion of the convolution kernel. Optimal performance is achieved at d = 2. The d = 1 represents the standard convolution. As the dilation rate increases beyond 2, we can see the performance start to decline.

### B. Spatio-modality attention for features attribution

In this experiment, we explore the feature’s (from all modalities) contribution toward the disease classification. The proposed spatio-modality attention inscribes these significance scores. Thus, for feature attribution, we refine the attention values. However, to summarize the scores, averaging over multiple dimensions might drastically suppress the data hence weakly depicting the tangible statistics. Here, we compute spatio-modality attention scores on a trained model by constraining multiple criteria. The values are summarized across the subjects and input feature dimensions. Furthermore, we compute scores feature-wise from all three modalities and determine a threshold representing the global mean throughout all the attention masks. We select the significant contributions by passing attention scores greater than the threshold only. Then, we filter the influential features by selecting significant contributions across at least 10% of the total population. The attention from all the settings mentioned in the reproducibility section is scrutinized to find a stable set of features. Therefore, we determine a contributing feature by their significant contribution toward classification across a reasonable number of samples. We avoid the features that are highly contributing to a few samples but are inconsistent across the population. As these sources are unstable and spurious, they are not reliable for neurological interpretation. Figure 4 (a) shows the sMRI feature analysis based on spatio-modality attention values. By thresholding the attention score, we observe four components to be significantly contributing to differentiating schizophrenia. These components are the caudate, supplementary motor area (SMA), temporal pole, and insula. These components are the caudate, supplementary motor area (SMA), temporal pole, and insula. Moreover, we examine their ICA loadings on both healthy control (HC) and schizophrenia (SZ) groups and visualize the group-level statistics in Figure 4(b). The HC loadings group mean is higher than the SZ mean, which indicates that HC subjects have higher gray matter density in the SMA region than SZ subjects. Alternatively, the SZ subjects are more heavily expressed in the insula and caudate than the HC group. Of note, the structural components differentiated through our method are associated with schizophrenia e.g., caudate [79, 80] and temporal pole [81]. Prior research also reported that aberrant motor behavior in schizophrenia is associated with supplementary motor area (SMA) volume [82, 83]. Also, insula activation has been associated with the processing of emotional facial expressions, which is deficient in individuals with SZ [84, 85]. The differences in gray matter density patterns between these groups aid in understanding the brain’s structural alterations associated with schizophrenia. We also run the two-sample t-test on the loading values from both subject groups (HC/SZ) to verify the statistical significance of the group differences. We found three of the salient features (Caudate, SMA, and Insula) show statistically substantial differences between control and individuals with schizophrenia at a level of *p* < 0.05. Furthermore, we analyze the attention computed from fMRI features. The static functional connections are shown in the connectograms of Figure 5. The connections are weighted by the attention scores in Figure 5 (a). The connections among sensorimotor (SM), cognitive control (CC), and subcortical (SC) are identified as most salient for classification. We further analyze the static functional connectivity of these edges between distinct brain components. Figure 5(b) shows average connectivity strengths are mostly negative (blue) and a few are positive (yellow). That demonstrates that most components are inversely correlated. The HC connectograms in Figure 5(c) show a connection with positive connectivity strength between IC 3 of subcortical to IC 10 of sensorimotor, whereas a positive connection in SZ dynamic is visible between IC 1 from SC to IC 30 of the cognitive control domain. The HC dynamics are more strongly connected in the subcortical and sensorimotor regions than the dynamics of SZ subjects. The average functional connectivity strength between the components of these two regions is higher for HC subjects than for SZ. The visual (VIS) domain also shows connectedness with other domains in HC dynamics where the SZ connections are comparatively weaker. The weaker connectivity in SZ might create functional impairments and dysfunction [86, 87]. Moreover, these connectivities also symbolize the communication between different parts of the brain for carrying out neural activities. Further study of these connections is required to assess their roles in overall cognition and information processing within these subject groups.

**Fig. 4.**
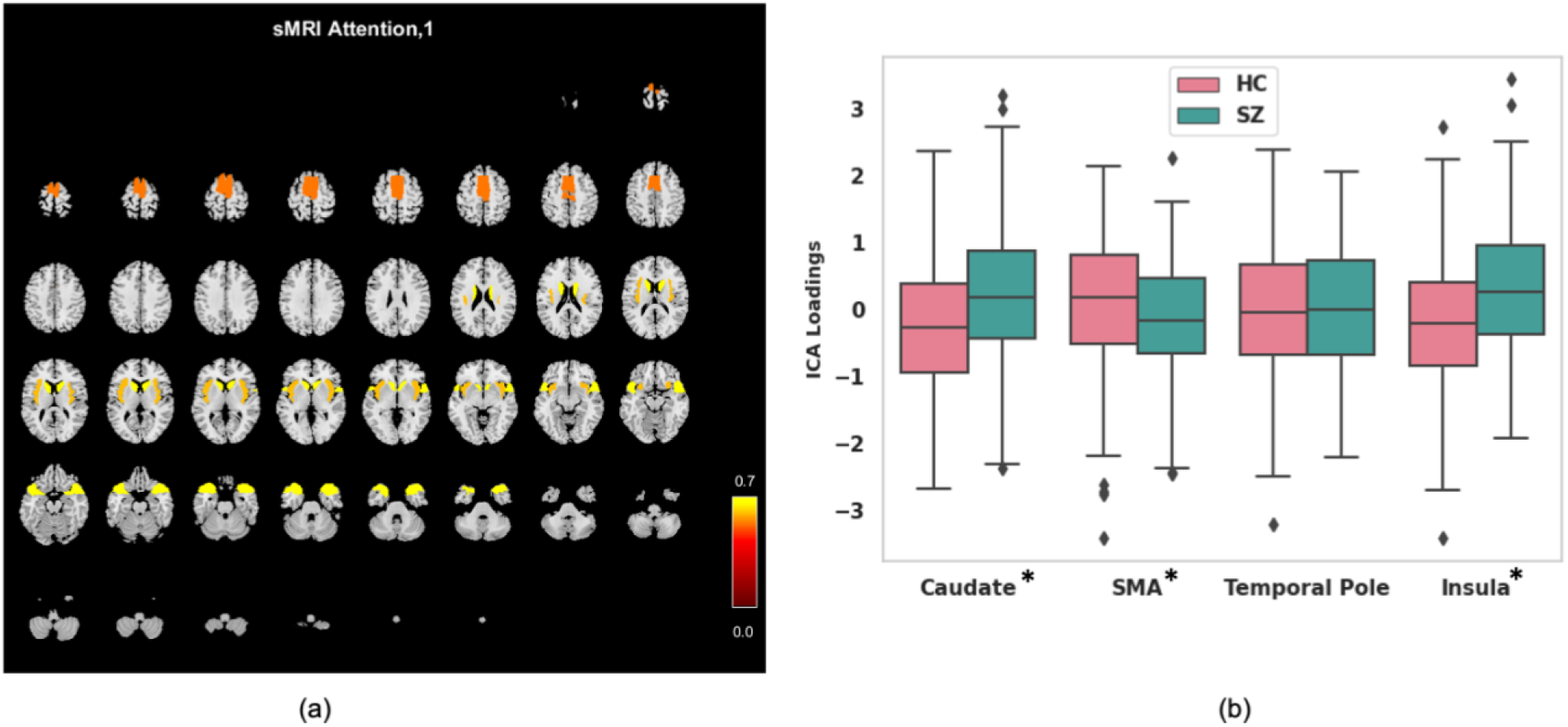
The spatio-modality attention-based sMRI features analysis. (a) Attention scores are computed on all sMRI components, and the significant ones are visualized on a structural montage. The components are supplementary motor area (SMA), caudate, temporal pole, and insula. (b) The ICA loadings for the most contributing components in HC and SZ groups. The error bars represent the standard deviation (SD). The asterisks sign on component name stand for statistical significance of the difference between HC and SZ. We run two-sample t-test to validate the differences. We use p-value < 0.05 to mark the significant differences.

**Fig. 5.**
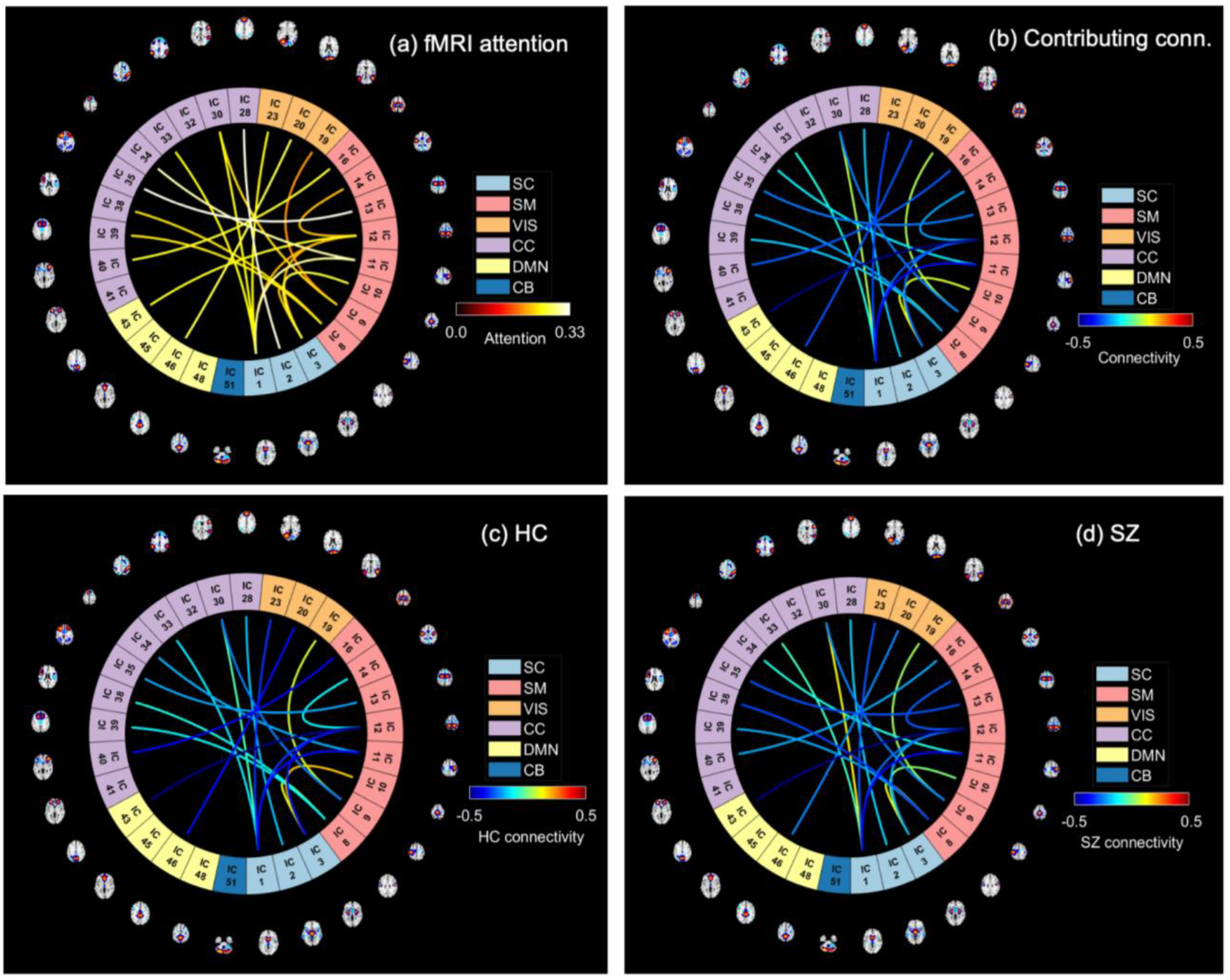
The spatio-modality attention scores computed from fMRI features (static functional connectivity (sFNC)). (a) The sFNC connections are weighted by their corresponding attention score. After thresholding, we show the connections that are effective for predicting schizophrenia at all different configurations of the model executations. The warm-colored connections are the significant ones; highly contributing in 5(a). We observe several contributing connections among sensorimotor (SM) and cognitive control (CC) and subcortical (SC) regions (b) The connections are weighted with the mean connectivity across the subjects group. (c) Connections are weighted by the mean connectivity strength computed across the HC group. (d) SZ group’s connectivity.

### C. Biclustering using spatio-modality attention scores

The spatio-modality attention masks provide the significance scores for the features from all modalities. In this experiment, we run clustering of these scores across modalities to examine the subgrouping of subjects and features based on their association with schizophrenia. Another rationale behind this experiment is to visualize the relation among distinct biological domains and their coregulation in diseased conditions. To run the analysis, firstly, we concatenate the feature’s attention from sMRI, fMRI, and genomics modality which is a two-dimensional matrix of (subject × features). Then, we run the N-BiC biclustering [88] for clustering the attention scores in both dimensions. We choose N-BiC because it allows clustering without specifying the expected number of biclusters and regulates the overlap between clusters. For two-dimensional data, it requires a heuristic to represent attributes from one variable as a function of attributes from another variable. It then exhaustively searches for all intrinsic subgroups and conditionally merges along the way. We sort features based on their consistently significant contributions to the classification. We select a subset of subjects for each feature where the attention values are higher than the global mean – the average attention across all the subjects and features. We choose features that exhibit this higher value across at least 10% of the total subjects. Initially, N-BiC yielded 4 biclusters then merged into 2 to regulate overlap. We repeat the run 10 times to stabilize the results. The outcomes show 1 multi-modal and 1 unimodal bicluster (Table 3). The multi-modal bicluster includes the following sMRI components: the anterior cingulate cortex (ACC), the medial prefrontal cortex, the caudate, the temporal pole, and the Frontal. It groups three SNPs including four genes CSMD1, ATK3, MOB4, HSPE1, and 26 static functional connections among several brain regions. AKT3 provides instructions for making a protein that is mostly active in the nervous system. The gene creates learning and memory-related deficits, and the SNP is identified as an associated genome-wide significant locus for schizophrenia [89, 90]. CSMD1 is known as a complement regulatory gene and has also been associated with schizophrenia [91, 92]. Figure 6 visualizes the sFNC connections through HC, SZ, and HC-SZ connectograms, respectively. In general, HC and SZ connections appear homogenous and rational since they are clustered into a single subgroup. However, one connection emerged with a strong HC-SZ difference between the cognitive control (IC 28) and sensorimotor network (IC 12) even with the maximized homogeneity. These multi-modal features analogously contributed to characterizing SZ; therefore, it potentially provides a link among these physiologies for disease-related deficits. These strong homogenous associations with schizophrenia suggest potential co-aberration of these physiologies (genomics and brain) suggesting that further exploration of their coupling and progression may help improve our understanding of the disorder. Further analysis of these features would be productive to infer their joint modulation in SZ conditions. Also, a tri-modal feature set can potentially help explain the behavioral deficits from multiple biological perspectives.

**Table 3.**
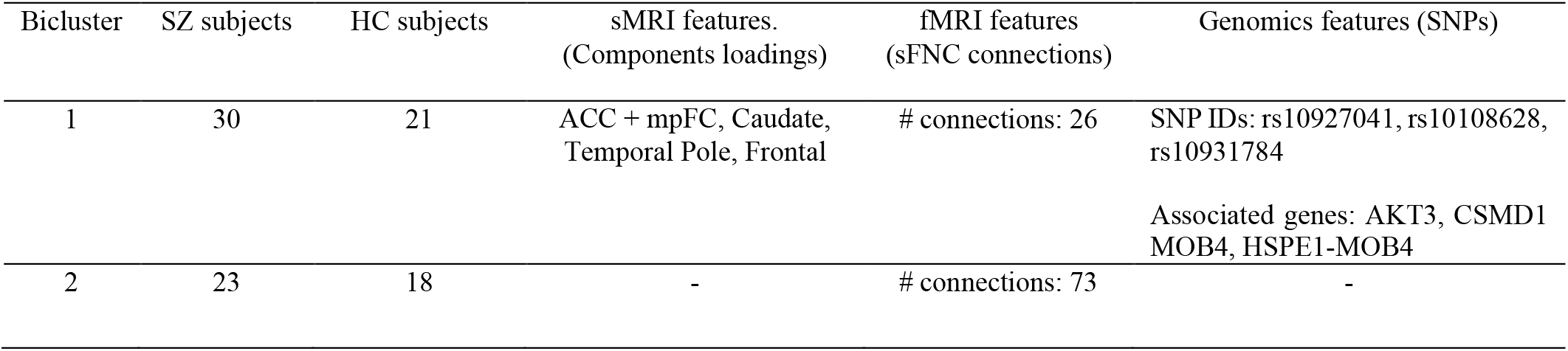
The clusters extracted from biclustering on the attention values from all three modalities. The N-BiC primarily extracts 4 biclusters that are merged into 2 due to higher overlaps with the earlier ones. Bicluster 1 is multi-modal and bicluster 2 is unimodal consisting of 73 sFNC connections.

| Bicluster | SZ subjects | HC subjects | sMRI features.<br>(Components loadings) | fMRI features<br>(sFNC connections) | Genomics features (SNPs) |
| --- | --- | --- | --- | --- | --- |
| 1 | 30 | 21 | ACC + mpFC, Caudate,<br>Temporal Pole, Frontal | # connections: 26 | SNP IDs: rs10927041, rs10108628,<br>rs10931784<br>Associated genes: AKT3, CSMD1<br>MOB4, HSPE1-MOB4 |
| 2 | 23 | 18 | - | # connections: 73 | - |

**Fig. 6.**
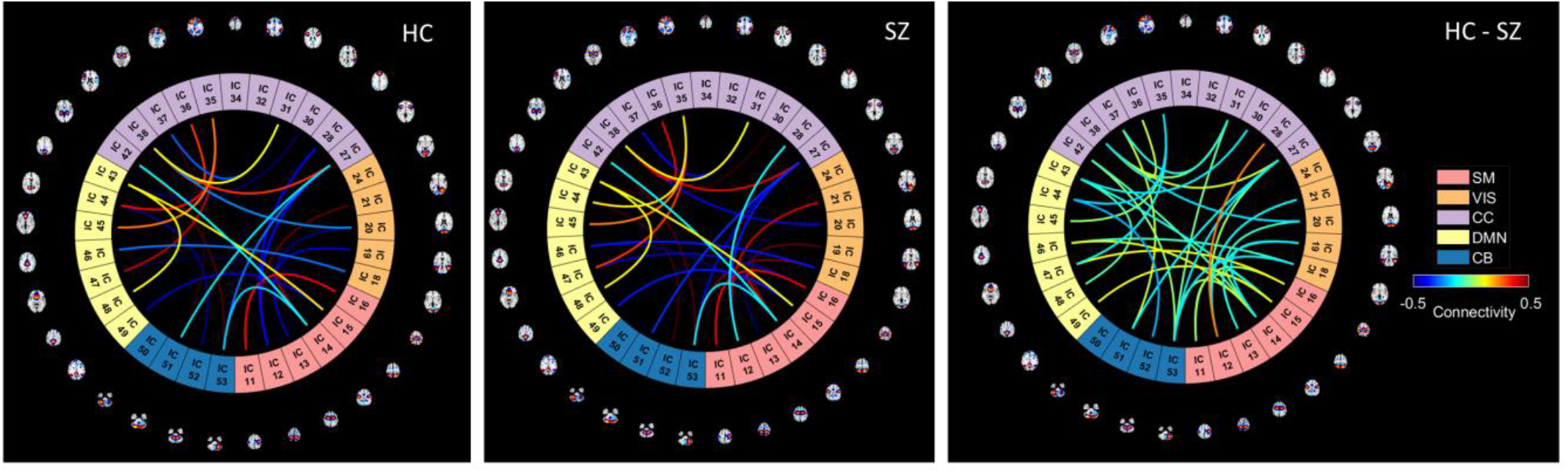
Connectograms generated from sFNC connections included in the multi-modal bicluster (bicluster 1). There are HC/SZ group differences in the strength of static functional connectivity in cognitive control (CO) to sensorimotor (SM), and visual (VS) regions.

## V. Conclusion

Our proposed fusion model attentively probes the dimensions of the fused latent space and provides a more rational synthesis of multiple biological sources. The spatio-modality attention module boosts the classification performance and achieves an accuracy of 94.1% with a 0.697 F1 score. The noteworthy performance of the classifier with spatio-modality attentive fusion evidences the utility of such a technique in multi-modal learning. Moreover, the spatio-modality attention scores are potentially self-explaining for the feature attributions towards the downstream task and provide an avenue for running further experiments e.g., subgrouping, and illustrating group differences. Our model learns spatial patterns through a large (dilated) receptive field for a better representation. The usage of dilated convolution for spatial scoring is potentially effective for foraging contextual information. In multi-modal settings, the dilation is shown to be providing a comprehensive view of the available modes of data. The modality submodule signifies each source and generates modality-wise contributions. As such, the controlled scoring also interprets the model’s decision and offers insights into neurobiological relevance. This relevance can potentially explain the underlying substrate of the disorder under investigation, schizophrenia. In all, the model seeks relevant patterns in the brain’s structural features, functional mechanisms, and genomic pathways that lead to a coherent deciphering of SZ. Additionally, the analysis of modality-specific contributions helps discriminate SZ-affected physiologies with limited availability. The subgrouping of subjects and multi-modal features based on the attention scores can potentially manifest the co-regulation of multiple biological domains in diseased conditions. The proposed fusion module can discover reliable biomarkers for the disorder, and the preceding interpretation recommends features that can help explain the underlying mechanism of the disease.

## Acknowledgments

The study was funded by National Institute of Health (NIH) grant number R01MH118695 and National Science Foundation (NSF) grant number 2112455.

## Author Contributions

Md Abdur Rahaman (MAR) and Yash Garg (YG) initialized using a bottleneck attention module for multi-modal data fusion. MAR and Vince D. Calhoun (VDC) designed the spatio-modality multi-source fusion applied on imaging genetics data. MAR ran the experiments and drafted the manuscript. Zening Fu (ZU), Armin Iraji (AI), and Jiayu Chen (JC) helped to preprocess the datasets and edited the manuscript. Peter Kochunov, Elliot Hong, Theo G. M. Van Erp, and Adrian Preda collaborated in data collection and edited the manuscript.VDC supervised all aspects of the project, thoroughly edited the paper, and gave valuable feedback on the results. All authors have approved the final version of the submission.

## Code and Data availability statement

Based on the IRB data privacy agreement, we are not allowed to share any subject-specific data. However, the datasets are available online in the referred studies and can be requested from the corresponding authors suggested in the studies. The preprocessing pipelines are public, as referred to in the manuscript. All the preprocessing tools are available online at http://trendscenter.org/software, and the model’s architectural details are available from the corresponding author.

